# The Cerebellum Implements an Oscillatory Forward Model for Accurate Motor Timing

**DOI:** 10.1101/2025.11.11.687835

**Authors:** Timothy O. West, Meaghan E. Spedden, George O’Neill, Sven Bestmann, Gareth Barnes, Simon F. Farmer, Hayriye Cagnan

## Abstract

Voluntary movements are composed of small sub-movements generated by pulsatile bursts of muscle activity at 4-10 Hz. These *motor intermittencies* synchronize with rhythmic brain activity and vary with movement velocity and sensory delays, suggesting they are a signature of a central oscillatory motor control mechanism. We hypothesized that these pulsatile movements arise from a cerebellar clocking mechanism-a neuronal timing process that flexibly adjusts its dynamics to maintain motor precision under temporal uncertainty. If so, cerebellar rhythms should modulate their influence on movement depending on cue predictability. To test this, we used optically pumped magnetoencephalography (OP-MEG), which enables high-quality, movement-tolerant recordings of the brain with dense cerebellar coverage. Six participants performed an auditory-paced finger flexion–extension task at approximately 1 Hz with either regular or irregular cue timing to vary predictability. All participants exhibited clear 4-10 Hz intermittencies in their kinematics, synchronized to neural activity in the cerebellum and the posterior parietal cortex. Our connectivity analysis revealed for the first time, that cerebellar activity at this frequency, predominantly reflects sensory feedback, but during regular, predictable cueing, the cerebellum shifts to exert a greater feedforward influence on movement. Moreover, cerebellar oscillations were most persistently phase-aligned with motor intermittent rhythms during accurately timed actions, an effect that was absent under irregular cueing. These findings support the idea that the cerebellum implements an oscillatory forward model for motor timing and provide novel evidence for its ability to adjusting its role in coordinating sensorimotor integration according to sensory predictability. Motor intermittencies may thus represent the output of a cerebellar timing process that underpins precise voluntary action.

## Introduction

At first glance, human movement appears smooth and continuous-yet closer inspection reveals motion that is composed of small discontinuous sub-movements occurring at 4-10 Hz (Woodworth, 1899). These *motor intermittencies* are seen in the electromyogram as alternating bursts of agonist and antagonist muscle activity (Vallbo & Wessberg, 1993). This activity resonates around spinal reflex circuitry (Wessberg & Vallbo, 1995) but is also widely synchronized across cortical and subcortical structures (Gross et al., 2002; Jerbi et al., 2007; McAuley & Marsden, 2000). The amplitude of motor intermittencies robustly correlates with movement speed (Hall et al., 2014; Jerbi et al., 2007; Pereira et al., 2017). The frequency of the intermittencies can be manipulated by artificially delaying sensory feedback (Miall, 1996), suggesting that they arise from circuits that compensate for sensorimotor delays. Furthermore, motor intermittencies persist even in the absence of visual feedback (Doeringer & Hogan, 1998) and contain components from both exogenous (i.e., sensory driven) and endogenous (i.e., centrally generated) motor processes (Susilaradeya et al., 2019). These findings suggest that motor intermittencies reflect a fundamentally rhythmic control process, through which motor commands are updated and corrected (Loram et al., 2006; Russell & Sternad, 2001).

This process can be viewed as a dual-oscillator system in which a low-frequency neuronal oscillator in the central motor system predicts future movements by aligning its timing with a peripheral oscillator formed between the spinal cord and muscles. We propose that the cerebellum implements this mechanism via a 4-10 Hz oscillator that synchronizes with a peripheral spinomuscular oscillator, working to continuously minimize error between the two. Under predictable conditions, this cerebellar “clock” operates in feedforward mode, using its rhythm to time motor commands. Under unpredictable conditions, a feedback mode dominates, and the cerebellar oscillator shifts its phase permissively to reduce mismatch with incoming sensory signals. This is conceptually equivalent to a Kalman gain, acting to dynamically reweight the influence of internal predictions versus sensory feedback upon upcoming motor plans (Diedrichsen et al., 2010).).

This accords with long held view that cerebellar circuitry and recurrent connectivity with the motor and sensory cortices make it well suited for predictive control (Hull, 2020; Ivry et al., 2002; Miall, 1996; Miall et al., 1993; Ramnani, 2006), and that motor intermittencies reflect dual processes of feedback and feedforward control (Susilaradeya et al., 2019). Recent work has shown that oscillatory cerebellar signals can build expectations of sensory inputs (Andersen & Lundqvist, 2019) in a way that is contextually sensitive to volatilities in timing (Andersen & Dalal, 2021). Furthermore, work shows that motor intermittencies synchronize not only with the cerebellum but also with the posterior parietal cortex (PPC), a region that is thought to maintain sensory predictions from the cerebellum for comparison with incoming feedback (Wolpert, Goodbody, et al., 1998). This suggests that both structures should be synchronized to peripheral motor intermittencies, and that their coupling should change under changes in the predictability of motor timing.

To test our hypothesis, we use an externally timed finger flexion-extension task with regular and irregular auditory cues to manipulate uncertainty in motor timing – a task shown previously shown to drive motor intermittencies synchronized with the cerebellum (Pollok et al., 2005, 2008). In this study, we have leveraged advances in optically pumped magnetoencephalography (OP-MEG; Boto et al., 2018) to provide a new window onto the cerebellar dynamics involved in motor timing of volitional movement. This new neuroimaging method provides improved robustness to movement artefact, and the absence of cryogenic cooling permits dense sensor coverage over the cerebellum. We have shown previously that this technology is able to localize putative cerebellar sources (C. Lin et al., 2019; C.-H. S. Lin et al., 2025), including those involved in pathological tremor (West et al., 2025). Using these tools, the present study looks for evidence of a phase-locked forward model of motor kinematics in the human cerebellum and tests predictions that its influence is sensitive to the temporal volatility of external cues. By demonstrating that cerebellar oscillations implement a phase-locked forward model, this work addresses a fundamental gap in our understanding of how the brain maintains temporal precision of movement.

## Results

### Intermittent Kinematics in Hand Movement and Associated Synchronized Neural Activity

Healthy human participants (n = 6) each performed on average 1972 ±135 repetitive, discrete finger flexion-extension movements on each hand, aiming to initiate flexion of all fingers (excluding the thumb) at the start of an auditory tone delivered with delays that were either regular (fixed delay of 800 ms) or irregular (random delay of 600, 800, or 1200 ms; Figure 1A). Each flexion-extension cycle was aimed to be completed before the start of the next cue. Participant 2 was found to perform the task in a continuous, sinusoidal tracking motion, and so movement-to-cue delays were not meaningful. For this reason, this participant’s data was excluded from the time-locked analyses (Section 2.4 onwards). A triaxial accelerometer was affixed to each subject’s index finger (Figure 1B) and brain activity was recorded using OP-MEG with sensor placements optimized to increase sensitivity to cerebellar and posterior cortical sources (Figure 1C-E). Recordings were divided into two-minute blocks of: (1) no movement; (2) movement-triggered by regularly paced auditory cues; or (3) movement-triggered by irregularly paced auditory cues (see Figure 1F).

**Figure 1.**
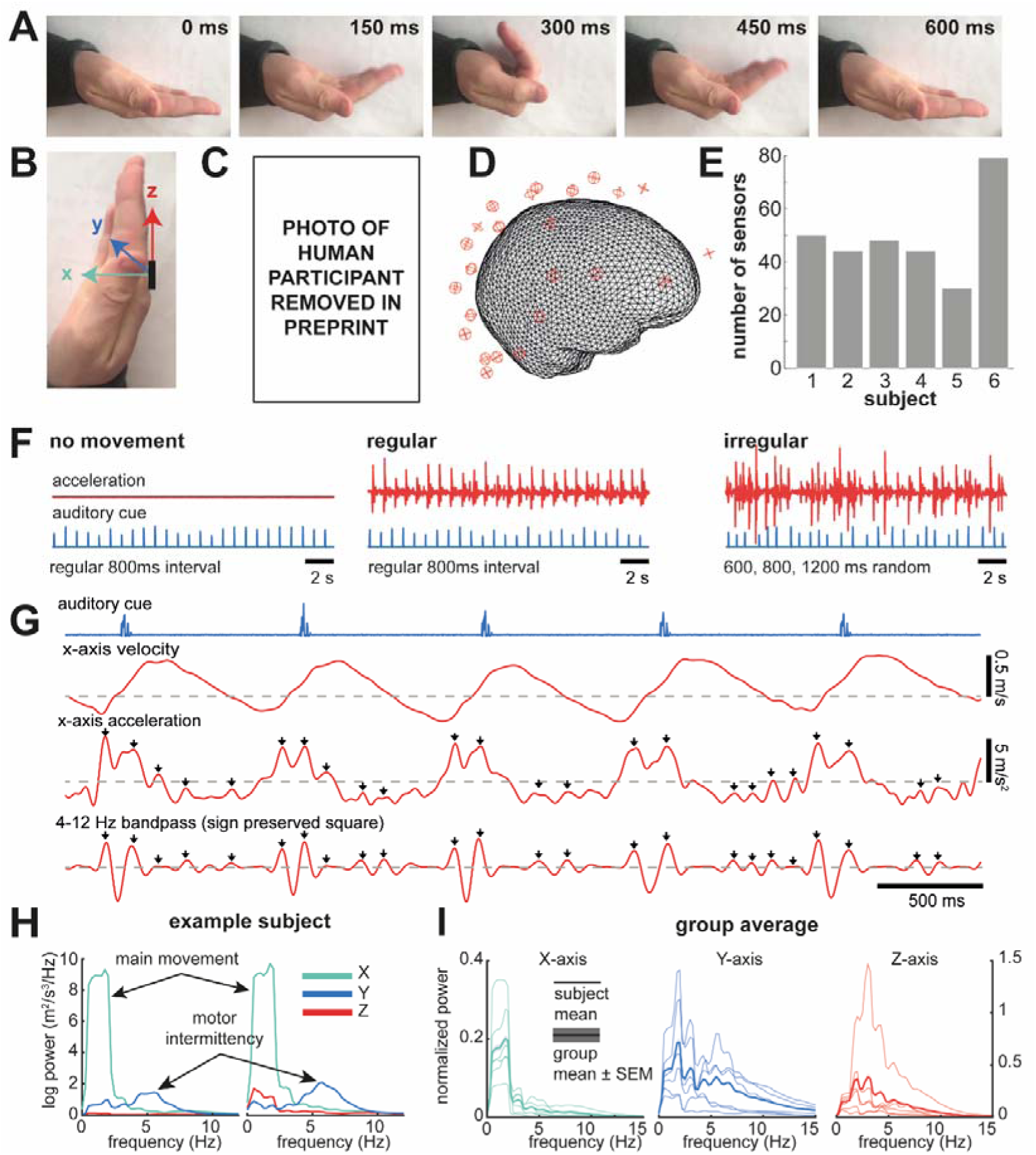
Optically Pumped Magnetoencephalography (OP-MEG) recordings of neural activity associated with human motor intermittencies in auditory cued finger movements. **(A)** Freeze frames from a video of a participant performing the cued finger flexion/extension task. **(B)** Schematic of orientations of accelerometer axes as mounted on the fingers. **(C)** OP-MEG sensors were mounted on a scanner-cast with cables suspended over the top front of the head. **(D)** Example of co-registration of source model with OP-MEG sensor layout. Sensors were positioned low on the neck to maximize sensitivity to potential cerebellar sources. **(E)** Total number of sensors per subject. **(F)** Subjects were asked to make auditory-cued movements at regular or irregular periods for both left and right hands, separately. A set of trials were also recorded in which the auditory cue was played but no movement was made. **(G)** Example traces over five finger flexions in response to an auditory cue (blue) shows clear motor intermittencies. Arrows indicate peaks in the broadband accelerometer signal corresponding to an underlying intermittency at 4-10 Hz (bottom trace). **(H)** Example subject’s power spectra of triaxial acceleration during movement trials. There is a clear peak at 0.5-2 Hz relating to the auditory paced movement, and a second peak centered at 6 Hz reflecting intermittent motor activity. **(I)** Normalized **g**roup level power spectra for each sensitive axis, individual subject shown with thin lines, whilst the group average is shown in bold.

Motor intermittencies in participants’ finger movements were visually apparent by eye and in the time-resolved accelerometry (Figure 1G), with a pattern of 4-5 cycles of intermittencies nested within each finger flexion movement, with power of intermittent oscillation peaking close to the maximum acceleration of the flexion movement. Power spectra of the acceleration (examples shown in Figure 1H and I) exhibit a large peak corresponding to the gross finger movement (0.8-1.6 Hz) along the predominant axis of motion (sagittal plane/X-axis), and a second smaller peak at 4-10 Hz corresponding to the motor intermittency.

The frequency and bandwidth of the intermittent frequency was stable across the six subjects (peak frequency: 5.4 ±0.75 Hz). Intermittency was most prominent in the transverse and frontal planes (Y- and Z-axes) showing on average 2.6 ± 3.0 times more power than in the primary movement axis (t(11) = 3.06, P = 0.011). These secondary axes contain less of the gross flexion–extension signal, making the intermittent component more distinct. For this reason, subsequent analyses primarily focus on these axes, where intermittency can be more reliably isolated. No difference in power for regular versus irregular cues was found (cluster permutation statistic of group level spectra).

The delay of participants’ movement responses to the auditory cues was 72 ms on average (one sample t-test, t(11) = -3.52, P = 0.005; Supplementary Figure 1). This is in line with the phenomena of negative asynchrony where subjects tend to act in anticipation of a cue (Aschersleben, 2002). Comparison between the distributions of delays between regular and irregular cueing shows that high cue volatility increases the likelihood of participants making early movements (cluster permutation statistic, -487 to -312 ms before cue, cluster t-value -2.79, P = 0.007).

### Motor intermittencies exhibit coherence with the ipsilateral cerebellum and posterior parietal cortex

To localize brain regions synchronized with kinematic intermittencies, we applied Dynamic Imaging of Coherent Sources (DICS; Gross et al., 2001) in 2 Hz wide frequency bands spanning 4-10 Hz. We compared the resulting kinematic-cerebral coherences between movement and data in which accelerometer trials were shuffled across trials, whilst the ordering of OP-MEG data was held constant (Figure 2). We selected the accelerometer axis that yielded the largest sensor level OP-MEG coherence as the reference signal for each subject. Maps of the overlap in peak t-statistics (|T| > 1.96) for each of the four frequency bands are shown in Figure 2. This analysis indicates that 8/12 hemispheres exhibit significant coherence in the contralateral posterior parietal cortex (PPC; blue arrows), an effect seen most strongly in the 4-6 Hz range. Coherence with putative cerebellar sources were found in 8/12 hemispheres and were most prominent at the high frequency range from 6-10 Hz. The peak in the overlap was localized to the contralateral cerebellum, lobule VIII (MNI: [-34, -56, - 52]), a region previously identified to exhibit BOLD activation during finger movement (Wiestler et al., 2011).

**Figure 2.**
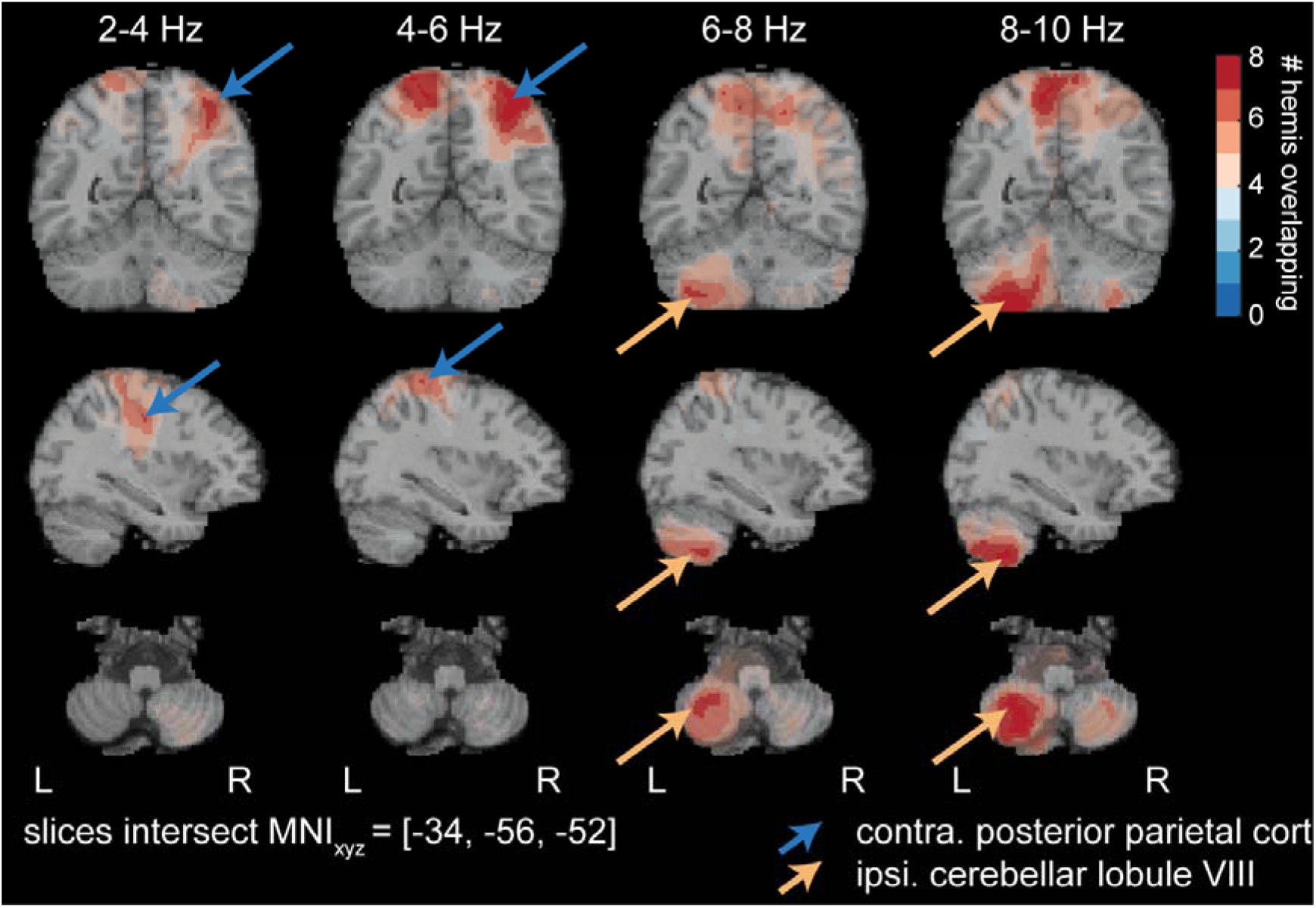
Imaging of sources synchronized between the brain and accelerometer at 4-10 Hz identifies key regions in the lateral cerebellum and posterior parietal cortex that synchronize to intermittencies during finger movements. Dynamic imaging of coherent sources (DICS) beamformer was used to identify sources in OP-MEG data synchronized to the motor intermittency. Coherence was computed between OP-MEG sources and the accelerometer and then compared against data in which the trial order of accelerometry data were shuffled, whilst the OP-MEG order was held constant. Images show the conjunction number between the resulting maps of T-statistics (|T| > 1.96) for movement in each hemisphere. Black and blue arrows indicate locations of ipsilateral cerebellar lobule VIII (MNI: [-34, -56, -52]) and contralateral posterior parietal cortical (MNI: [36, -62, 56]) sources that exhibit increased synchronization to motor intermittencies during movement. Maps were computed using four common filters using data filtered at 2-4 Hz, 4-6 Hz, 6-8 Hz, and 8-10 Hz. A complementary source map, with slices centered on the posterior parietal cortex can be found in Supplementary Figure 3.

Note due to sensor placement at the back of the head, frontal sources such as premotor cortex were not resolved in this analysis (see Methods). A secondary analysis of source power at 4-10 Hz is shown in Supplementary Figure 2. These results indicate that intermittencies in finger kinematics are synchronized with a parietal-cerebellar circuit during movement. We next examine how these changes are expressed in the frequency domain and assess the direction of neural connectivity within and between the periphery and cerebrum.

### Cerebellar-Kinematic Coherence Predominantly Represents Peripheral Feedback

To better understand how the intermittent dynamics of hand movements are functionally related to neural activity within the cerebellum, we performed a spectral analysis of signals from virtual electrodes constructed within the ipsilateral motor cerebellum. Analysis of spectral power of the acceleration (Figure 3A) identified an increase in 4-10 Hz power during finger movements that was also reflected in increased power of the cerebellar virtual channel in this band during movement (Figure 3B, 1.5-8 Hz, cluster t-value 4.21; P = 0.003). Coherence analysis shows that within the same 4-10 Hz band there is strong synchronization between the ipsilateral cerebellum and peripheral accelerometry that is significantly increased in movement relative to rest in 6/6 participants (Supplementary Figure 4) and significant at the group level (Figure 3C; 0.5-5.75. Hz; cluster t-value 2.36; P = 0.014). Interestingly, we also saw evidence of resting state beta frequency cortico-kinematic coherence that was significantly increased when the participants were not moving (cluster permutation statistic, 15.8-21.0 Hz; cluster t-value -2.71; P = 0.006), that may reflect a high frequency rest tremor.

**Figure 3.**
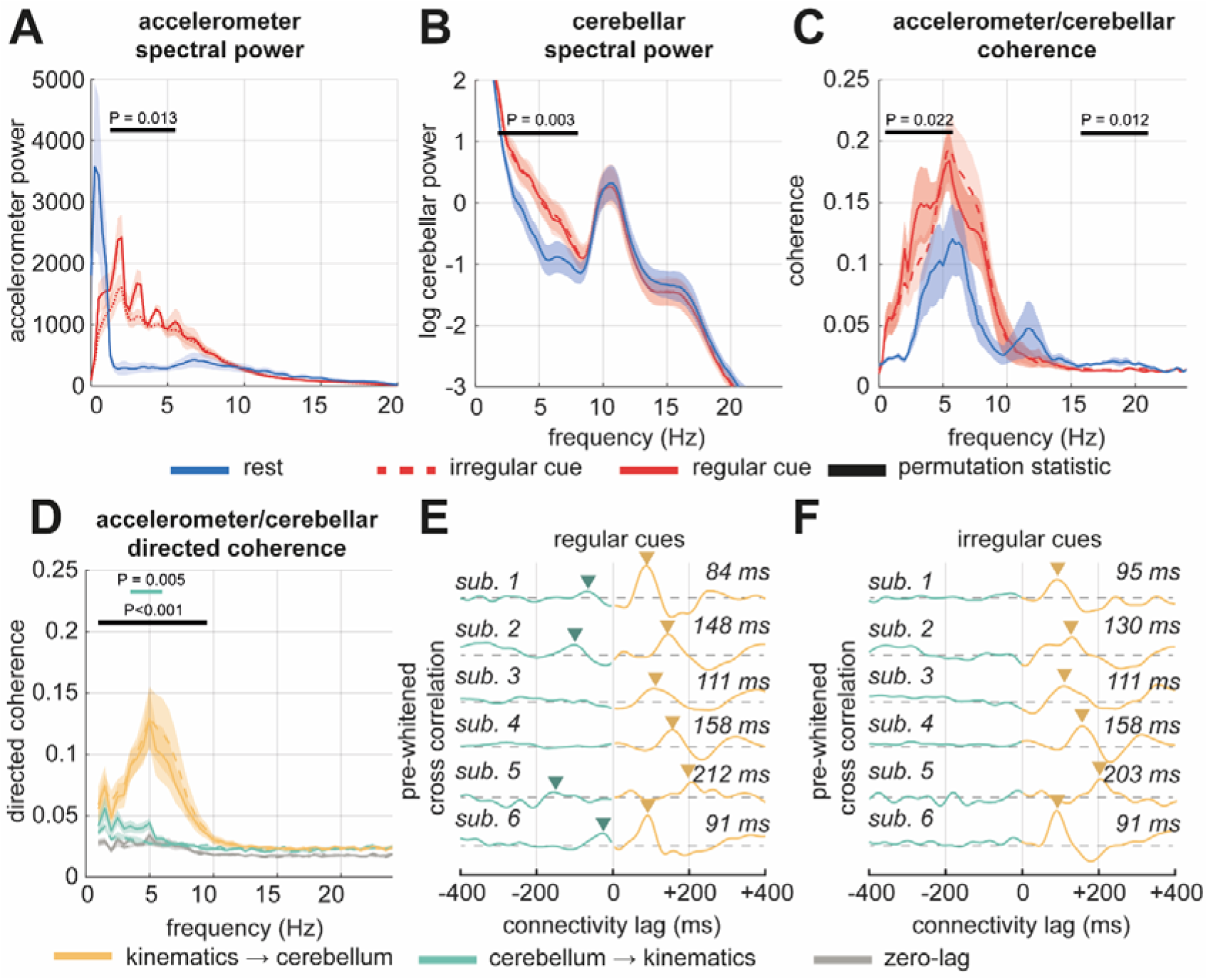
The cerebellum is strongly synchronized with motor intermittencies in the kinematics with lagged connectivity indicating a predominantly afferent pathway. Group averaged analyses of power spectra and connectivity were made using virtual channels constructed from the ipsilateral cerebellum and compared to the axis of the accelerometer with peak sensor average coherence. Left and right movements were combined by always taking the cerebellar source ipsilateral to the movement. Bars indicate significant permutation cluster corrected t-statistic for rest vs movement (black); regular vs irregular cues (in the color of its matching line). **(A)** Group averaged power spectra of accelerometer signal between movement (red) and rest (blue). Motor intermittencies show clear increase during movement. **(B)** Same as (A) but for OP-MEG virtual electrode projected to the ipsilateral cerebellum. Increases in neural activity at the intermittent frequency during movement are apparent. **(C)** Group averaged coherence spectra indicating synchronization between finger acceleration and cerebellar neural activity. During movement, coherence between these signals is strengthened within the intermittent band. **(D)** Directional influences were estimated using non-parametric directionality. Group averaged spectra are shown for zero-lag (grey), cerebellum to kinematics (efferent; green), and kinematics to cerebellum (afferent; yellow). Bars indicate significant permutation cluster corrected t-statistic for afferent vs efferent directionality (black); regular versus irregular cues for forward and reverse directions (red and blue, respectively). **(E and F)** The pre-whitened time-domain cross-correlation indicates lagged influence of kinematic intermittencies upon the cerebellum with an average delay ∼150 ms, for regular and irregular cue conditions. Corresponding subject level analyses of power and coherence/NPD are shown in Supplementary Figures 3 and 4, respectively.

Having established the existence of significant cerebellar-kinematic coherence, we next decomposed the coherence into its directional components using non-parametric directionality (NPD), a robust measure of directed functional connectivity (Halliday, 2015; West et al., 2020). All subjects showed a significant coherence peak in afferent connectivity (i.e., accelerometry to the ipsilateral cerebellum) in the 4-10 Hz band (Figure 3D), that was significantly stronger than the reverse efferent connection (i.e., ipsilateral cerebellum to the accelerometer reference axis; cluster-permutation statistics, 3.0-9.0 Hz; cluster t-value -3.95; P <0.001). Examination of the pre-whitened cross-correlation (Figure 3E and F) revealed clear peaks for 6/6 participants corresponding to a cerebellar delay of +157 ±44 ms relative to the accelerometer reference.

To test our hypothesis that predictable cues increase cerebellar influence on kinematic intermittencies, we compared NPD spectra between trials with regular and irregular movement cues (bold and dashed red lines, respectively). This analysis revealed a significant increase in efferent connectivity from cerebellum to intermittent limb kinetics in the 4-5 Hz range for the regular cue condition (4/6 participants, 3.5-6.0 Hz; cluster t-value 2.86; P = 0.005), with peaks visible in the associated pre-whitened cross-correlation (Figure 3E) A supplementary analysis of signals in the cPPC also found significant afferent coupling with the accelerometer that increased during rest, but we found no evidence for modulation of coupling by cue regularity in this region (Supplementary Figure 4).

Evidence for a functional connection between the cerebellum and PPC at the intermittent frequency was also found using NPD (Supplementary Figure 5), with 4/6 subjects showing significantly larger connectivity in the cerebellum to PPC direction (P < 0.05). These data suggest that intermittencies predominantly reflect a feedback signal in the cerebellum and posterior parietal cortex, likely integrating sensory information from the periphery. However, in line with our hypothesis, when cue timings become more predictable, a conditional feedforward component driven by the cerebellum provides greater influence upon intermittent finger kinematics.

### Intermittencies Synchronize Across a Cascade Startin from the Periphery to Motor Cerebellum and Posterior Parietal Cortex

The localization of synchronized activity to kinematic intermittencies in the cerebellum and PPC (Figure 2), and the existence of significant afferent directional coherence between these structures and the periphery (Figure 3), accords well with their involvement in motor control. To test our hypothesis that motor control circuits exhibit time dependent synchronization to motor intermittencies, we investigated oscillatory dynamics using both the amplitude envelope and the phase locking value (PLV; Lachaux et al., 1999) – a measure of time varying phase synchronization - at the frequency of the kinematic intermittency (Figure 4). To do this, we used the beamformer to construct virtual electrodes placed in the ipsilateral cerebellum and contralateral PPC. To ensure phase estimates were stable, we used a narrowband FIR bandpass filter at 5 ±2.0 Hz based upon the group averaged coherence between the cerebellum/PPC and accelerometery on the hand (Figure 3).

**Figure 4.**
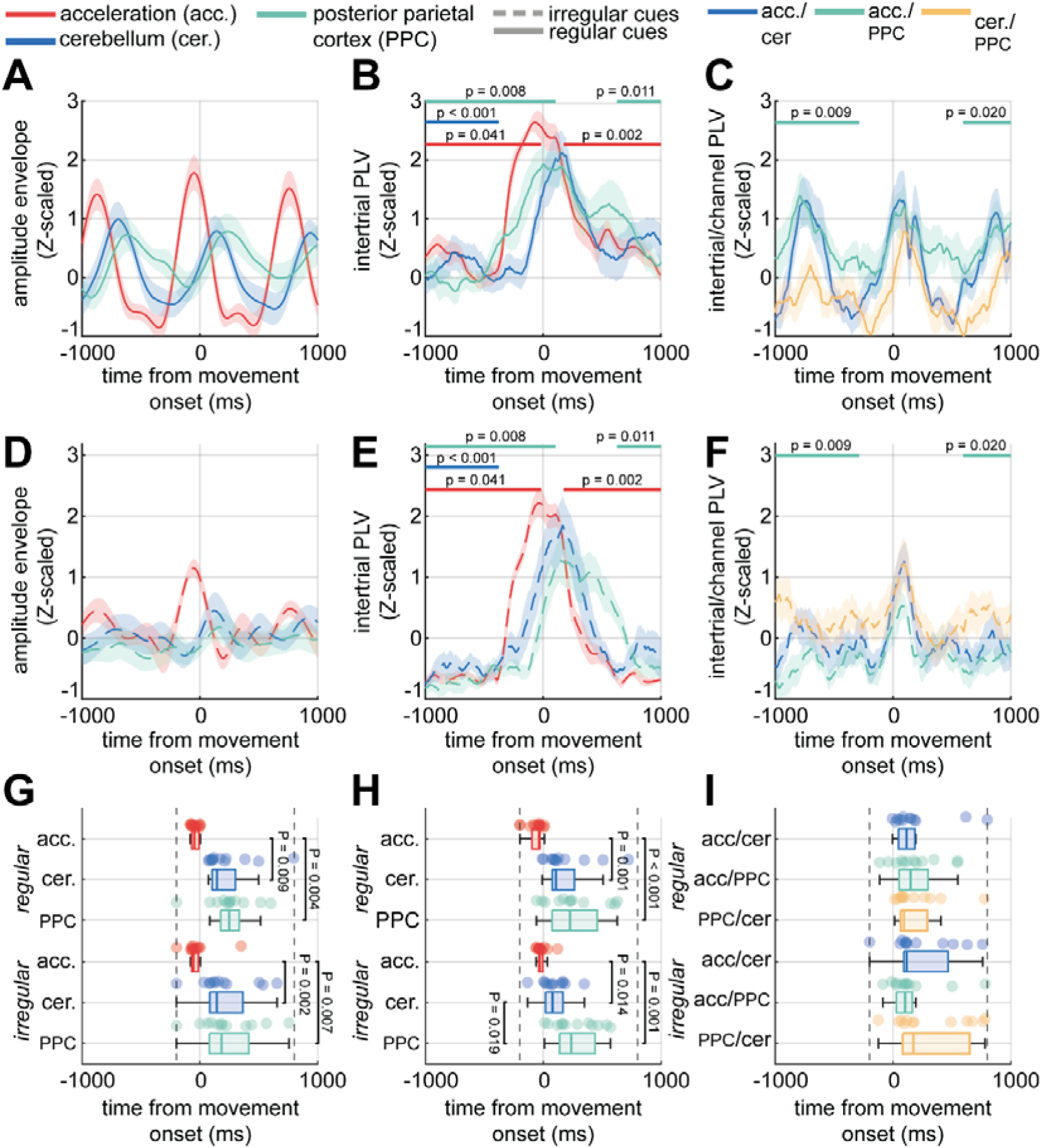
Time locking analysis of oscillatory synchronization shows cerebellar and parietal signals cascade from motor intermittencies in the periphery. Signals from accelerometry and OP-MEG virtual channels in the cerebellum and posterior parietal cortex (PPC) were time-locked to the peak acceleration of each finger movement (i.e., t = 0). Analyses are bandlimited to 4 ±1.5 Hz. **(A)** Group averaged amplitude envelopes of the subject specific motor intermittency band for intermittent acceleration (i.e., accelerometer axis with most power in 4-10 Hz band; red), the motor cerebellum (blue) and the PPC (green). Bars indicate a significant cluster permutation statistic between regular and irregular cues. **(B and C)** Same as A, but for the between trial between-trial phase locking value (PLV, i.e., across trial phase alignment), and between trial, between channel PLV. **(D, E and F)** Same as A, B, and C, but for irregularly paced auditory cues **(G)** Box plots of time of amplitude peak. Bars indicate P-values of a two-sample t-test exceeding α = 0.05. Dashed lines indicate the range for which peaks were defined. **(H and I)** Same as G, but for the between-trial, between-channel PLV (i.e., trial consistency of phase differences), for pairs: Accelerometry/Cerebellum (blue), Accelerometry/PPC (red), Cerebellum/PPC (green). A corresponding analysis performed when time locking to auditory movement cues is presented in Supplementary Figure 6.

Analysis of the amplitude envelope dynamics during regular and irregular cued movement (Figure 4A and B) reveals that the intermittencies detectable in cerebellar and PPC activity reflect a cascade of activity arising from an afferent process that originates in the periphery. There is a clear peak in amplitude of motor intermittencies on average 45 ms before movement onset. Analysis of the timing of peaks in the amplitude envelopes (Figure 4G) demonstrates that brain activity at the motor intermittent frequency occurs significantly later, with a peak of +282ms in the PPC (regular cues; paired t-statistic(11) = -4.47, P = 0.009) and +293 ms in the cerebellum (regular cues; paired t-statistic(11) = -4.95, P = 004). No significant difference in onset times between the cerebellum and PPC were found. These data suggest a clear cascade of intermittent activity from periphery to the PPC and cerebellum during finger movements.

We next analysed the intertrial PLV to assess trial-by-trial phase consistency (Figure 4B and C). High PLV values reflect a strong phase reset of the intermittent rhythm around movement onset (t = 0 ms). In both regular and irregular cue conditions, signals related to motor intermittencies showed a period of phase reset on and around movement onset. However, overall intertrial synchronization was stronger under regular, predictable cueing. Specifically, we observed significant PLV increases in the accelerometry both before and after movement onset (-1000 to -15 ms and 76 to 1000 ms; cluster t-value > 4.0; P = 0.002), in the cerebellum before movement onset (-1000 to -373 ms; cluster t-value 2.36; P = 0.026), and in the PPC before and after movement onset (-1000 to 107 ms and 627 to 1000 ms; cluster t-value > 3.0; P < 0.008). Together, these results show that predictable cueing enhances the stability of intermittent phase dynamics across the periphery, cerebellum, and PPC during finger movement.

Timings of intertrial phase synchrony (Figure 4H) followed a similar sequence to amplitude envelopes, with both motor cerebellar and PPC peaks delayed from the accelerometery (PPC: regular cues; paired t-statistic(11) = -4.51, P < 0.001; Cerebellum: regular cues; paired t-statistic(11) = -4.20, P = 0.001). Again, no differences in timing were found for the cerebellar and PPC peak phase locking. Thus, the phase resetting in cerebellar and parietal regions follows on from kinematics in the periphery.

Both cerebellum and PPC also exhibit significant interregional phase synchronization (Figure 4C and F), although no differences in onset timing was found for this analysis. Neither the cerebellum nor PPC exhibited strong phase reset in response to the onset of auditory cues (Supplementary Figure 6). These data suggest that the circuit between the periphery, the motor cerebellum and posterior parietal cortex exhibit a cascade of phase synchronization at 4-10 Hz, starting in the muscles and then propagating to the cerebellum and PPC around similar times. The raised level of phase synchrony between different movements supports our hypothesis that intermittency provides the substrate for a continuous clock signal that keeps persistent timing during highly predictable timing conditions.

### Accurate Motor Timing is Associated with Enhanced Cerebellar Phase Locking

In this final section, we directly test our hypothesis that accurate motor timing is facilitated via phase-locked synchronization associated with increased phase stability of a cerebellar oscillator operating at the same frequency as motor intermittencies. To examine this, we analysed the trial-by-trial variations in instantaneous phase and looked for intertrial phase resets followed by extended periods of phase consistency related to the degree of timing error. This analysis is presented as a raster plot showing the phase evolutions of cerebellar and kinematics rhythms across all trials recorded in the experiment, with rows sorted by the movement delay relative to cue, as a measure of timing error (Figure 5).

**Figure 5.**
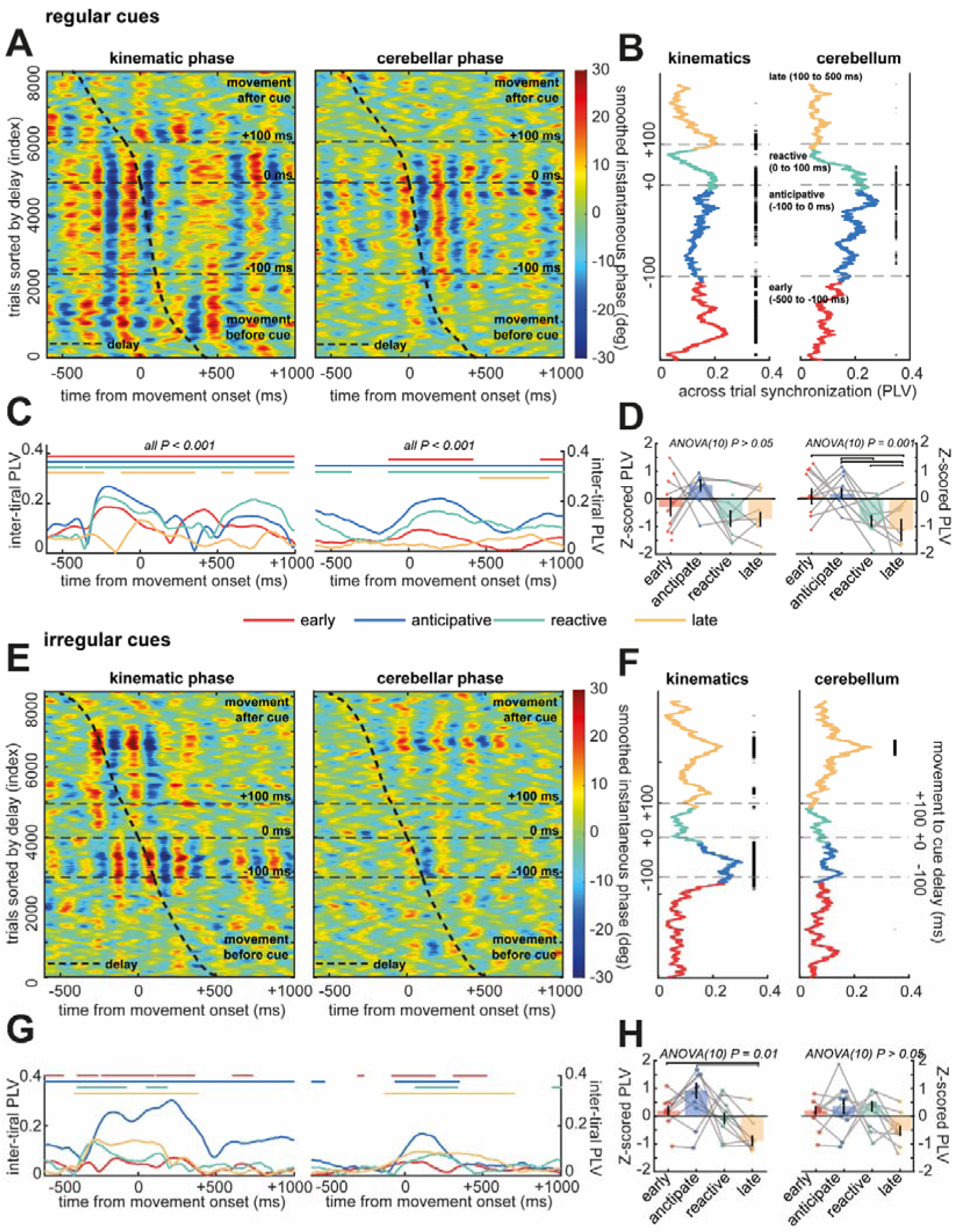
Phase dynamics of motor intermittencies in kinematics and the cerebellum are organized by movement timing. Instantaneous phase was extracted on a trial-by-trial basis from accelerometery and cerebellar virtual channels, time-locked to peak finger acceleration (t = 0). Trials were categorized by movement timing relative to cue into four groups: early (–500 to –100 ms), anticipatory (–100 to 0 ms), reactive (0 to 100 ms), and late (100 to 500 ms). Group level analysis shown. Panels A to D, and E to H, indicate analysis of regular and irregular cues, respectively. **(A)** Raster plot showing instantaneous phase across all trials, sorted by motor delay relative to cue (indicated by the thick dashed line). Consistent phase bands reflect inter-trial phase locking. **(B)** Inter-trial phase locking value (PLV) plotted against movement delay. Black bars indicate significant phase locking compared to phase-randomized surrogates (P < 0.001). Peak cerebellar PLV aligns with trials showing minimal delay (|delay| ≈ 0). **(C)** Time-resolved PLV profiles stratified by timing category. Bars indicate significant test against the phase randomized surrogate (P < 0.001). Kinematic phase locking precedes movement and presents similarly across trials. Cerebellar phase locking increased following movement onset and was strongest in more accurate trials. **(D)** Subject-level analysis of the PLV–delay relationship (as in panel B) shows significant modulation (ANOVA) by movement delay for cerebellar, but not kinematic, signals. Points and lines indicate the profiles for each left and right movement set (i.e., two hemispheres). Example of a single subject analysis is shown in Supplementary Figure 7. **(E-H)** Same as A-D, but for irregularly cued movements.

**Figure 6.**
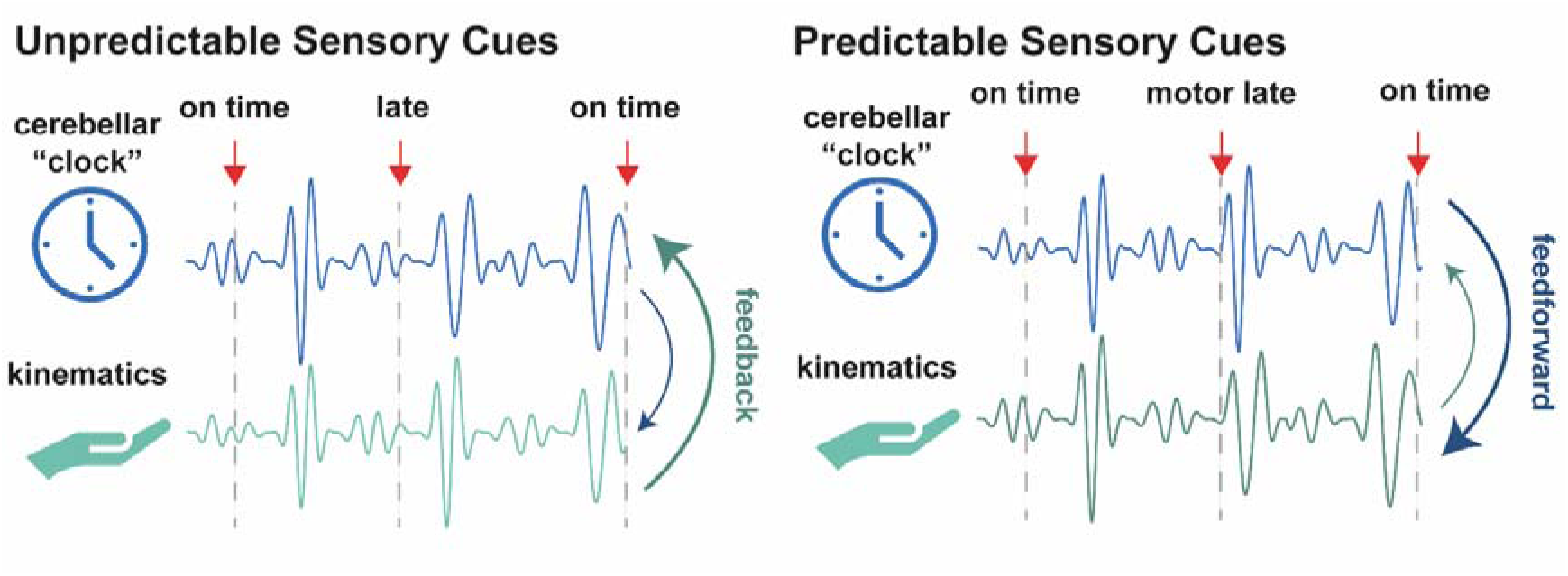
Our analyses of neural dynamics in the cerebellum are consistent with an internal clocking mechanism for the accurate timing of finger movements. This paper has tested the hypothesis that cerebellar dynamics synchronize to motor intermittencies and provide an oscillatory forward model that can facilitate accurate motor timing. Our analyses of directed functional connectivity shows that cerebellar synchronization to 4-10 Hz kinematic intermittencies are predominantly representative of afferent connectivity, however, when cues become highly regular, the cerebellum exhibits increased influence in the efferent dynamics – a feature consistent with increased feedforward control of movement. This mechanism is also consistent with our analysis of instantaneous phase by which accurate motor timing is associated with more persistent and stable cerebellar oscillation.

Our analyses of recordings made during predictable, regularly cued movement show that there is significant phase consistency within kinematic intermittencies (Figure 5A, left panel) that exhibited significant intertrial phase synchronization (measured as the intertrial PLV) across wide periods of delays ranging from -400 to 150 ms relative to the auditory cue (Figure 5B, left panel; comparison to surrogate distribution, bars indicate P < 0.001). This synchrony remained high throughout the whole movement (Figure 5C left; comparison to surrogate distribution, bars indicate P < 0.001). Across hemispheres the degree of phase locking in the kinematics was not associated with timing accuracy (Figure 5D left; ANOVA (10), P > 0.05).

In contrast, when the same analysis was applied to cerebellar activity, we found that phase locking is strongest in trials when movement timing was accurate (Figure 5A, right panel), with the intertrial synchronization peaking for trials with less than 75 ms error in timing (Figure 5B, right panel; comparison to surrogate distribution, bars indicate P < 0.001). The timing of phase locking for accurately timed movements is also clustered around 150 ms after movement onset (Figure 5C, right pnael) with timing of the peak shifting for very pre-emptive (100 ms before the cue; in red) or delayed (greater than 100 ms after the cue; in yellow) movement trials. This modulation of cerebellar phase locking was significant across hemispheres at the group level (Figure 5D; ANOVA (10), P = 0.001) with anticipatory movements (-100 to 0 ms delay) showing significantly greater intertrial PLV than for those that were very late (in yellow). These group level analyses were representative of that seen at the single subject level (Supplementary Figure 7).

During irregular, unpredictable motor cueing, phase synchronization of kinematic intermittencies was present and again occurred across wide ranges of motor delays (Figure 5E and F, left panels). There was evidence that phase synchronization to intermittencies was reduced between very early and late response (Figure 5H left panel; ANOVA(10), P = 0.01). However, when analysing activity in the cerebellum, there was a marked loss in overall phase synchronization (Figure 5E and F, right panels) comparing to that analysed during regular cueing. As expected, this was accompanied by a loss of delay specific modulation in cerebellar phase locking (Figure 5 H, right panel; ANOVA(10), P > 0.05).

These findings provide strong evidence that accurately timed movements are associated with increased phase stability of a cerebellar oscillator. This oscillator becomes stable and thus synchronized across trials in which there is high temporal precision, and in trials were cues were temporally regular. This supports the hypothesis that oscillatory activity in the cerebellum instantiates a dynamic, synchronized control mechanism for timing motor output.

## Discussion

### Summary of Results

Using OP-MEG during auditory-cued finger movements, we identified robust 4-10 Hz kinematic intermittencies that were consistently exhibited across participants (Figure 1) significantly synchronized with neural activity localized to the ipsilateral cerebellum and contralateral posterior parietal cortex (PPC; Figure 2). During regular, isochronous movements, directional connectivity analyses indicated that, while afferent feedback dominated overall, cerebellar efferent influence on limb kinematics increased significantly (Figure 3), suggesting stronger feedforward control when timing was predictable. Phase-locking analyses further showed that intermittent rhythms propagated from the periphery to cerebellum and PPC with delays of ∼150–250 ms (Figure 4). Crucially, cerebellar oscillations exhibited enhanced phase stability during accurately timed movements, with intertrial synchronization peaking around 75 ms after movement onset (Figure 5). These findings suggest that under predictable conditions the cerebellum stabilizes its intrinsic oscillator, strengthening its role as a forward model to guide motor timing.

By contrast, during irregular cueing, cerebellar and PPC synchronization remained predominantly afferent, with no increase in cerebellar outflow (Figure 3). Although motor intermittencies were still evident in the periphery, their neural correlates showed weaker phase stability and reduced trial-by-trial synchronization. The relationship between cerebellar phase locking and timing error was abolished (Figure 5). This indicates that under temporal uncertainty, the cerebellum shifts into a feedback-dominated mode, updating its oscillator more flexibly to incoming sensory signals rather than driving feedforward predictions. Together, these findings support a framework in which the cerebellum provides an oscillatory forward model for the accurate timing of motor control.

### Motor Intermittencies are Synchronized Across an Oscillatory Control Network

Our study builds upon a large body of work suggesting that intermittencies, rather than representing sensorimotor noise, are a direct signature of a cortico-cerebellar control process for fine movement. Previous work has shown that intermittencies result from paired flexor/extensor bursts that provide discrete updates of movement via alternating periods of acceleration and deceleration (Vallbo & Wessberg, 1993). Such alternating periods are synchronized with cerebellar and cortical activity (Gross et al., 2002; Jerbi et al., 2007). Our analysis of directed functional connectivity aligns with previous measures that suggest cerebellar-kinematic coherence is predominantly afferent– reflecting a process of sensory feedback. Both cross-correlation and time resolved analyses here suggest a delay of ∼150 ms that is larger than expected from spino-cerebellar transmission (∼30 ms), but comparable with previous time-locked analyses (Andersen & Dalal, 2021; Tesche & Karhu, 2000). These longer delays are consistent with the activity of a cerebellar oscillator that integrates information over multiple cycles of intermittency, rather than resets arising from singular events of spino-cerebellar transmission.

We report for the first time here that cerebellar outflow to the periphery increases in highly predictable sensory contexts such as that during isochronous regular cueing. This is line with previous reports of increased cerebellar-cortical coupling during accurate motor timing (Pollok et al., 2008). We speculate that this process implements an effective Kalman gain on the motor controller (Diedrichsen et al., 2010), with modulation of the cerebellar output via the deep cerebellar nuclei acting to shift between feedback processing during periods of uncertainty, and feedforward process when movement timing is predictable.

Synchronization of the PPC to motor intermittencies has been previously reported (Gross et al., 2002; Jerbi et al., 2007; McAuley & Marsden, 2000), but its role is unclear. Our finding of delayed PPC synchronization (∼150-250 ms post-movement) supports its role as a comparator region in forward model architectures (Wolpert et al., 1998), potentially receiving delayed input from cerebellar sources. This idea is also supported by our finding directed connectivity between these structures was predominantly led by the cortex, although we found no clear differences in timing of synchronization between the PCC and cerebellum.

### Cerebellar Phase Entrainment as the Neural Substrate of State Estimation

Persistent low frequency (< 10 Hz) oscillations have been repeatedly observed in inferior olive, the cerebellum’s main afferent nucleus (Welsh et al., 1995). Similar to that reported here, these oscillations are persistent across time (Hartmann & Bower, 1998) and strongly phase-locked to movement onset (Welsh et al., 1995). Similar oscillations in humans, whilst more difficult to detect, have been implicated in the accurate temporal prediction of somatosensory stimuli (Andersen & Dalal, 2021; Tesche & Karhu, 2000). Our findings extend this work by showing that the synchronization of cerebellar oscillations is also tightly associated with accurate movement timing. Our results show that phase is dynamically entrained, depending on the movement and the timing relative to cue – consistent with the cerebellar oscillator acting as a state estimator for movement.

These findings are consistent with the view that the cerebellum functions as a forward model within a discrete-time predictive control loop (Miall et al., 1993; Wolpert et al., 1998). Specifically, we propose that model is implemented in the phase dynamics of intrinsic cerebellar oscillations, which regulate the timing of efferent output through alignment with motor intermittencies. This model can explain the seemingly dual intrinsic/extrinsic nature of kinematic intermittencies (Susilaradeya et al., 2019). Here, we observed unstable cerebellar phase dynamics during inaccurately timed movements, suggesting that the cerebellum continuously updates its internal oscillatory model in response to timing errors. This aligns with ideas that cerebellar circuits minimize sensory prediction error through internal simulations of expected dynamics (Friston & Herreros, 2016). Understanding the function of healthy oscillatory cerebellar activity is relevant for pathology, in which dysregulation of healthy oscillatory control strategies may lead to pathological dynamics such as that seen in essential tremor (Steina et al., 2025; West et al., 2025) or ataxia (Martino et al., 2014).

### Limitations

Detecting cerebellar activity non-invasively has long been difficult due the structure’s deep location and convoluted neuroanatomy. However, the isolated mammalian cerebellum generates a noninvasively measurable field (Okada et al., 1987) and modelling work estimates cerebellar signals are attenuated by only 30–60% compared to cortical sources (Samuelsson et al., 2020). MEG studies have now consistently detected cerebellar activity in motor and cognitive tasks (Andersen et al., 2020). OP-MEG has been used to detect and localise cerebellar activity in an eyeblink conditioning task (C. Lin et al., 2019; C.-H. S. Lin et al., 2025) and in studies of pathological tremor (West et al., 2025). Our localization of activity to lobule VIII aligns with prior work that expects motor related activation in this area (King et al., 2019).

Our continuous, cue-driven task limited the ability to disentangle cerebellar contributions to anticipation, planning, and error correction. Future paradigms should incorporate preparatory periods or catch trials to better separate feedforward from feedback mechanisms such as that in (Andersen & Dalal, 2021). Advances in OP-MEG now enable studies with larger samples and more complex behaviours such as reaching and walking (O’Neill et al., 2023; Seymour et al., 2021; West et al., 2025), that can explore the generalization of the findings presented here. Moreover, whole-head sensor arrays will provide coverage of frontal and premotor regions-absent in the present study - allowing a fuller account of networks including frontal and premotor regions.

### Conclusions

This study provides the first direct evidence that the human cerebellum exhibits time-locked phase synchrony that is consistent with a dynamic forward model for motor timing. Using OP-MEG, we demonstrate that these cerebellar rhythms are phase-locked to kinematic sub-movements and modulated by the predictability of sensory input - supporting a phase-locked loop architecture for cerebellar control. Our findings suggest that motor intermittencies are not merely biomechanical artifacts but signatures of an active control process integrating afferent feedback and efferent prediction. By revealing how cerebellar oscillations stabilize during accurate movement, we offer new insight into the neural implementation of internal timing mechanisms and lay the groundwork for future exploration of cerebellar-cortical control across more naturalistic and complex behaviours.

## Methods

### Recruitment and Details of Participants

Ethical approvals for research with healthy participants was obtained from the ethics committees at University College London, with subjects’ consent obtained according to the Declaration of Helsinki. Six healthy participants were recruited for this study.

### Optically Pumped Magnetometer Recordings

Optically pumped magnetoencephalography (OPM) recordings were made using QuSpin-manufactured sensors mounted in 3D-printed rigid scanner casts (Chalk Studios, London, UK) custom-built to each participant’s scalp shape as determined using a structural head/brain MRI. 5/6 participants used 2^nd^ generation (dual-axis sensitive) sensors exclusively, whilst subject six used a combination of both 2^nd^ and 3^rd^ (tri-axial) sensors. In the absence of an MRI (subject 1), the head shape was estimated using an infrared depth camera. Offsets of 1-3 mm were added to scanner casts to allow for anatomical error, tissue deformation, and hair. One subject (subject 3) used an existing headcast designed for a similar headshape – comparison of scalp shapes showed <5% deviation. Experiments were conducted in a magnetically shielded room (MSR; Magnetic Shields Ltd, Staplehurst, UK). The inner layer of mu-metal lining the room was degaussed using a low-frequency decaying sinusoid driven through cables within the walls before the start of the experiment. The OPM sensors were then nulled using onboard nulling coils and calibrated. The OPM acquisition system (National Instruments, Texas, USA) had a sampling frequency of 6 KHz and a 16-bit resolution. An antialiasing 500 Hz low-pass filter (60th order FIR filter combined with a Kaiser window) was applied before data were down-sampled offline to 2 kHz. Task triggers were sent to the OPM DAC using a LabJack U3 DAC (LabJack Corp., Colorado, USA). Acceleration was recorded via a sensor affixed to the knuckle (Lilypad, Analog Devices Massachusetts, USA with custom made cable).

To render electromagnetic artefacts arising from movement of magnetometer cabling amenable to artefact correction algorithms, we bundled cables tightly with foam padding to minimize independent cable vibration. Cables were also positioned so that they passed over the front of the head to avoid interference with sensors close to cerebellum (Figure 1A). Sensors were placed preferentially at the back of the head to maximise sensitivity to cerebellar sources (Figure 1B). OPM recordings were made with 49 ±6 sensitive axes (Figure 1C).

### Movement Task

The motor paradigm involved a series of auditory cued whole finger flexion/extension of either the left or right hands. Participants were instructed to initiate a finger flexion at the start of each auditory cue, requiring temporal prediction of the cue start. The auditory cue consisted of a 200ms, 5 KHz tone. Cues were delivered with either regular (800 ms) or irregular (600, 800, or 1200 ms) intervals (figure 1D). Each flexion-extension cycle was aimed to be completed before the start of the next cue. Stimuli were delivered using scripts written with PsychToolbox-3 (Kleiner et al., 2007). Stimuli were played via ear tubes (except subject 5, cues played with speaker). The auditory waveform was simultaneously recorded using the auxiliary inputs of the OPM acquisition system. Data was also collected when the subject was at rest, with eyes open and listening to the regular set of auditory cues but not moving. There were 10 repeats of each of the 6 conditions, with each block lasting 2 minutes, performed for left and right movements separately.

### Preprocessing, Artefact Removal, and Data Rejection

Data were analysed using custom scripts written in MATLAB (The Mathworks, Massachusetts, USA) using the Fieldtrip (Oostenveld et al., 2011) and SPM 25 toolboxes (Tierney et al., 2025). Data were digitized and down sampled to 512 Hz. Individual OPM channels were visually inspected to remove those affected by gross movement artefact, and then high and low pass filtered at 2 and 98 Hz using FIR windowed-sync filters. Line noise was removed using a discrete Fourier transform – fitting sine and cosine components to the line frequency and then subtracting from the data. Homogenous Field Correction was applied to OPM data to achieve interference rejection (Tierney et al., 2021). Triaxial accelerometer signals were low passed filtered at 20 Hz using a using a FIR windowed-sync filter. Trials with remaining artefacts were rejected visually, using thresholds on channel maximum and average Z-scores, and kurtosis measures.

### Source Space Analysis and Statistics

Forward models for source space analysis used a single shell head model built from each participants’ structural MRI. In the case of subject 1, no MRI was available and so a template head model was used and rescaled using ICP to match an optical 3D scan of the participant’s head. A source model was constructed from a 3D grid with 1 cm spacing, applying a nonlinear warping to allow comparison of anatomy at the group level. Sensor locations were aligned to the subject’s MRI using ICP between the scalp segmented from the MRI and that used for the construction of subject specific headcasts. Source orientations were optimized for maximal power.

Dynamic Imaging of Coherent Sources (DICS) was used to map the distribution of coherence across the cerebral volume. Covariance matrices were computed for the motor intermittency band (4-12 Hz). We constructed beamformer weights for a common filter covering both movement and rest periods, and for left and right movements separately. A second wideband (2-98 Hz) beamformer was used to project sensor-level data to source-level virtual electrodes. Covariance matrices were truncated by their effective rank (Westner et al., 2022) and a covariance regularization of 1% was used. We term coherence between inertial and OPM signals cortico-kinematic coherence, following Bourguignon et al., (2011). DICS images were computed for 40 second blocks (consisting of ten trials of 4 seconds each).

Source space statistics were computed using SPM12. DICS images from right hand movements were flipped along the sagittal axis, so that the left hemisphere was consistently ipsilateral to movement. At the single subject level, a simple two-sample t-test was used to contrast movement with data in which the accelerometery trials were shuffled. Due to the small sample size available, we present overlap maps indicating the number of subjects expressing a |T| > 1.96 for a given voxel.

### Virtual Electrode Construction, Parcellation, Dimensionality Reduction and Orthogonalization

Using the wideband spatial filter described, we projected “virtual electrode” signals across a 1 cm spaced grid covering the brain. Each region of interest was assigned a brain parcellation based on the Automated Anatomical Labelling atlas 3 (Rolls et al., 2020). We reduced the dimension of the virtual electrode data by taking the 1^st^ principal component of signals contained within each parcel but preserving hemispheres. We constructed cerebellar ROIs using known motor regions(lobules I-IV, V, and VI) in line with known functional mapping (King et al., 2019). To mitigate source leakage arising in MEG data, we applied symmetric orthogonalization to these parcellated principal components prior to further analyses (Colclough et al., 2015).

### Spectral Estimates and Functional Connectivity Analysis

Spectral estimates utilized multitaper estimates with taper numbers set to achieve a 6 Hz (for source space analyses) or 2 Hz (for spectral plots) smoothing window. Functional connectivity was estimated using the Neurospec toolbox. Symmetric measures were estimated using magnitude squared coherence (Halliday & Rosenberg, 2000), whilst non-parametric directionality (NPD) was used to estimate directed connections (Halliday, 2015; West et al., 2020). Estimates were computed over 40 second blocks (consisting of ten trials of 4 seconds each). Statistics to compare between conditions or directions were achieved using cluster-permutation tests (cluster forming threshold at P = 0.1; 1000 repetitions).

### Time-locked Analysis of Movement and Cues

To investigate neural activity that was responsive to the initiation and changes in movement, we time-locked virtual electrode signals to limb kinematics. The X-axis (the principal movement axis) of the accelerometer signal was Z-standardized individually for each epoch of finger movement, and then a threshold at Z = 1. Breaks in threshold crossings less than 100ms were conjoined, and the first time point of each suprathreshold event used as trigger to time lock analysis of movement. Cue onsets were determined from the audio trace by taking Z = 2 as the threshold.

### Phase Synchronization Analyses

We used the phase locking value (PLV; Lachaux et al., 1999) to investigate the phase synchronization of neural and kinematic signals at the frequency of kinematic intermittency. We looked at both inter-trial phase coherence of single channels (i.e., a measure of consistent event related phase resetting), as well as intertrial, between channel phase coherence (i.e., a measure of consistent event related relative phase difference), and PLV computed within a sliding window (400 ms, 95% overlap). All PLV traces were converted to t-statistics using a baseline at -500 to -100 ms. Peak times of the PLV dynamics were computed by taking the maximum of each subject and hemisphere. PLV variance estimates were achieved by bootstrapping trials using 500 resamples with replacement. Phase estimates used for PLV calculations were computed by bandpassing (512 order windowed sync FIR) at 5±1.5Hz. Phase estimates were then made by taking the inverse tangent of the Hilbert transformed analytic signal. Intertrial phase raster plots were constructed by smoothing phase estimates across trials (50 trial moving average) and also time (50 ms moving average window).

Amplitude response curves were calculated using the pairwise phase differences for each trial, taking the circular average over the range dictated by the group statistics of the peak synchrony taken as the mean ± 1 standard deviation. The kinematic movement power for each trial was normalized as a percentage difference from the mean power. Confidence interval for the ARC was constructed from a surrogate distribution in which the trials’ phase and kinematic power were in a randomized order. Prior to group averaging, all ARCs were recentred to the maximum bin.

