## Supplementary Figures for "The Cerebellum Implements an Oscillatory Forward Model for Accurate Motor Timing"

### 1.1 Supplementary Figure 1

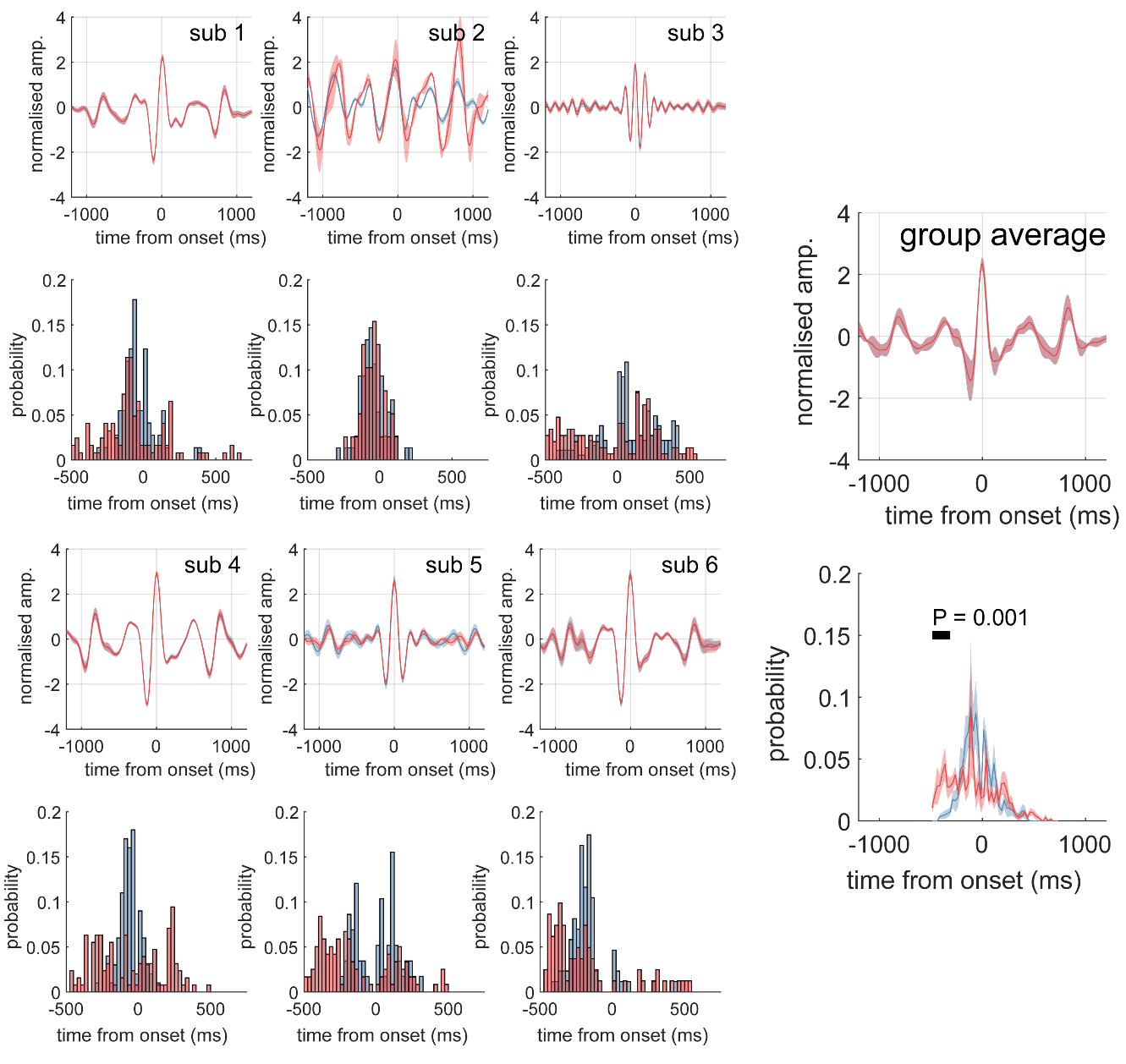

Supplementary Figure 1 – **Averaged responses of the finger movements across subjects.** For each subject the time averaged accelerometry trace is shown (main axis X) for regular (red) and irregular (blue) cues. The corresponding histograms show the distribution of movement delays relative to the cue onset times. The group averaged profiles (right) indicate a propensity for early responses in the irregularly cued movements.

### 1.2 Supplementary Figure 2

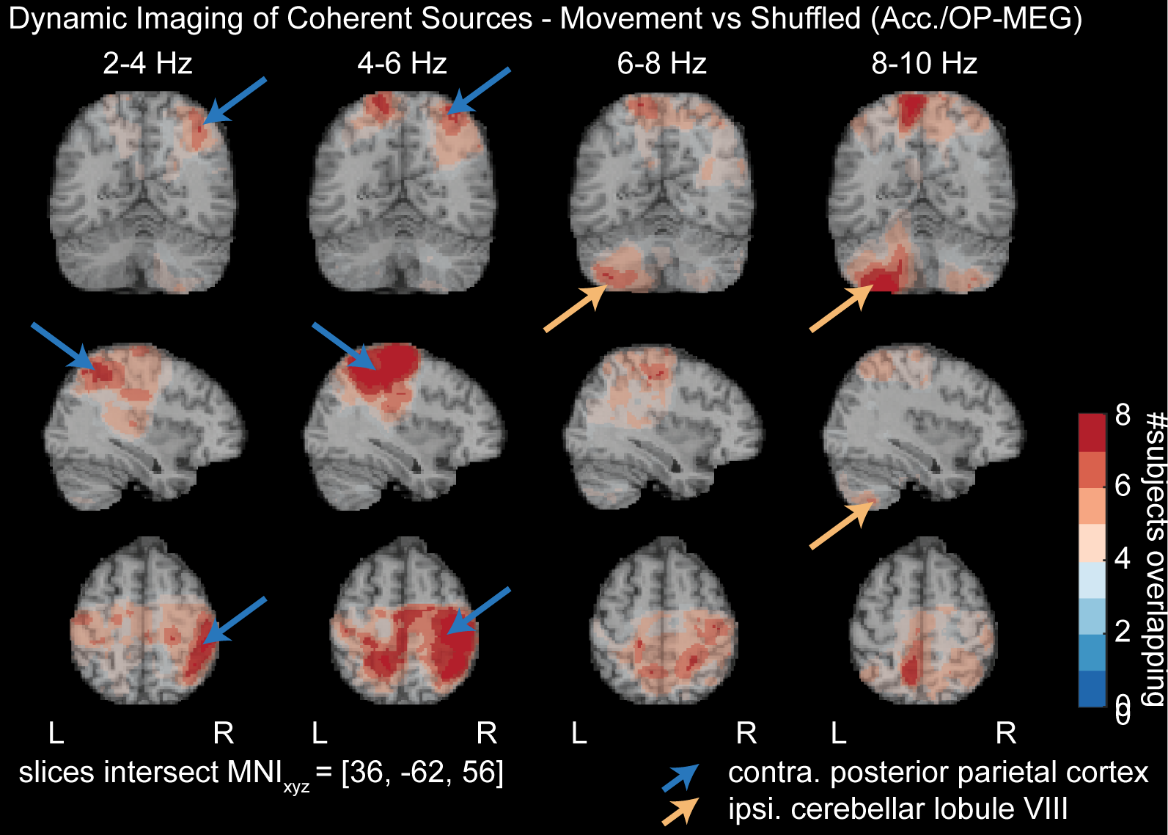

Supplementary Figure 3 - **Summary of group level ANOVA statistics comparing source localized OP-MEG power at the frequency of kinematic intermittencies at 4-10 Hz between regular vs irregular auditory cueing conditions.**

### 1.3 Supplementary Figure 3

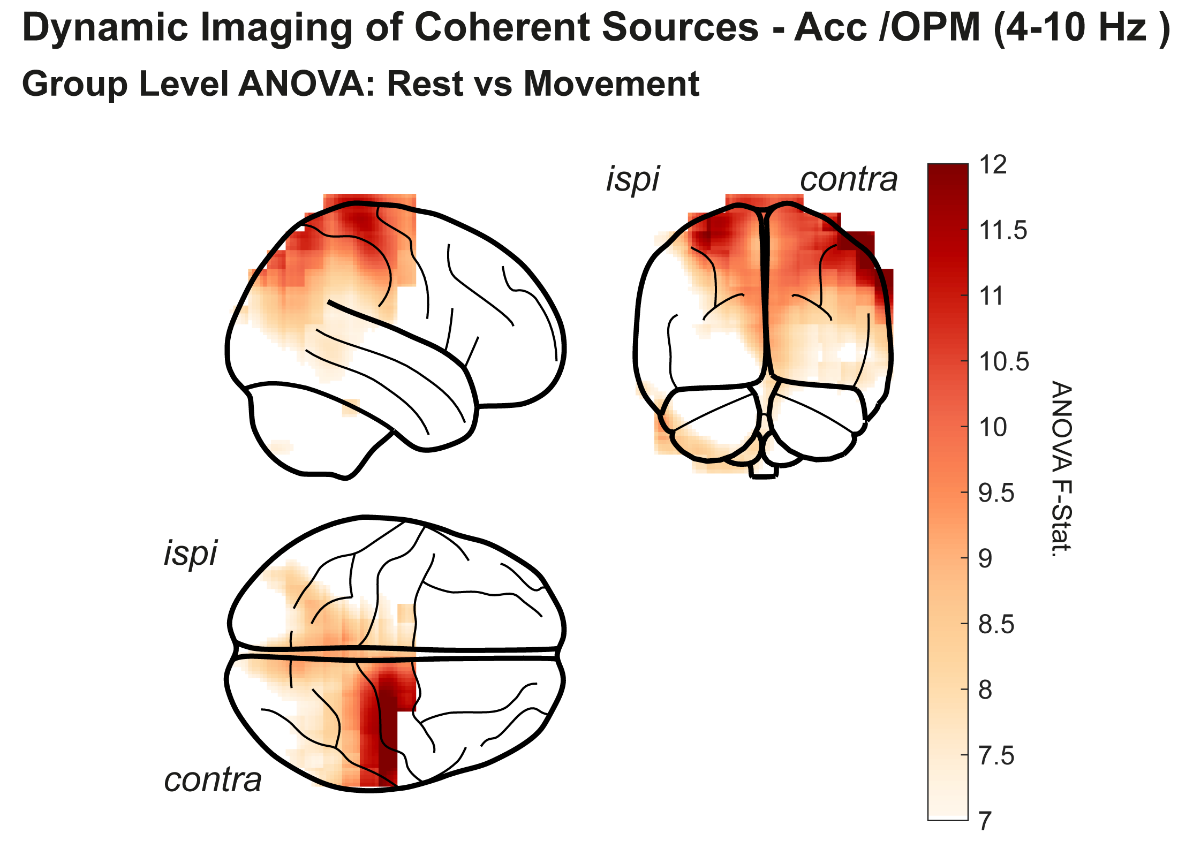

Supplementary Figure 3 - **Summary of group level ANOVA statistics comparing source localized OP-MEG power at the frequency of kinematic intermittencies at 4-10 Hz between regular vs irregular auditory cueing conditions.**

### 1.4 Supplementary Figure 3

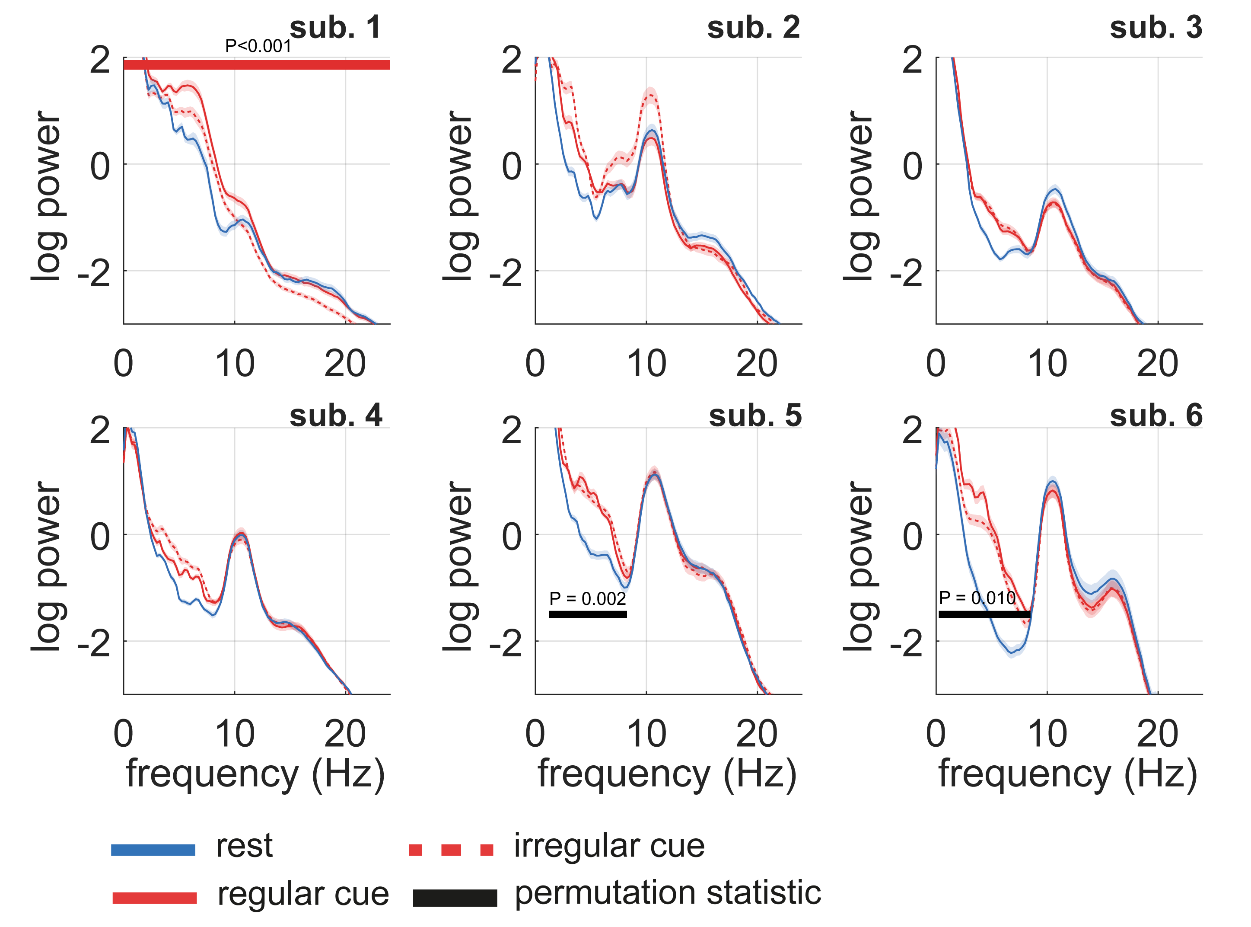

Supplementary Figure 3 - **Summary of changes in cerebellar power.** Power analyses were made using virtual channels constructed from the ipsilateral cerebellum. Line plots show trial/subject mean, and shaded region gives the standard error of the mean. Log scaled power spectra comparing movement (blue) and rest (red) showed for each of the five subjects. Bars indicate significant permutation cluster corrected t-statistic for rest vs movement (black).

### 1.5 Supplementary Figure 4

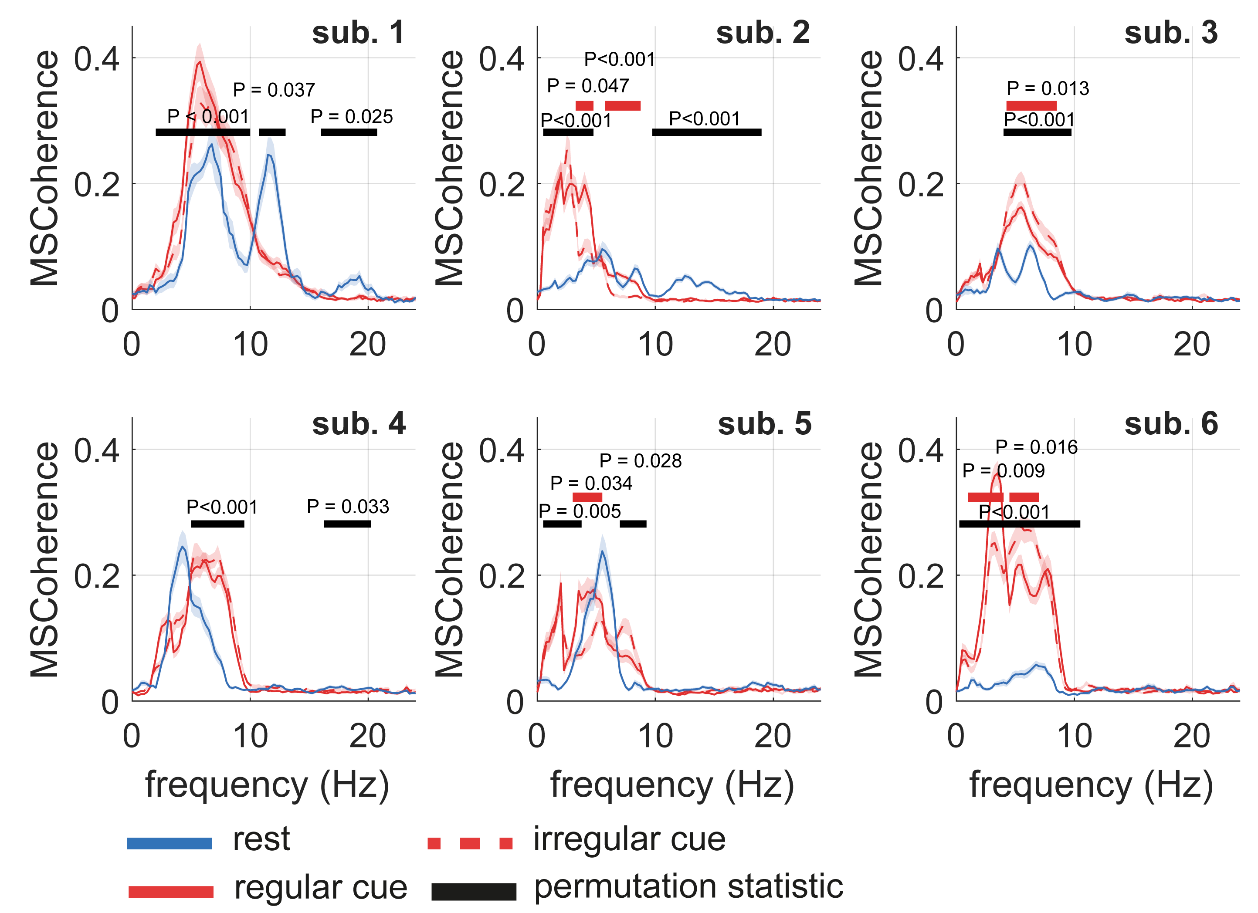

Supplementary Figure 4 - **Summary of changes in cerebellar/accelerometer coherence.** Coherence analyses were made using virtual channels constructed from the ipsilateral cerebellum. Line plots show trial/subject mean, and shaded region gives the standard error of the mean. Coherence spectra comparing movement (blue) and rest (red) showed for each of the six subjects. Bars indicate significant permutation cluster corrected t-statistic for rest vs movement (black).

### 1.6 Supplementary Figure 5

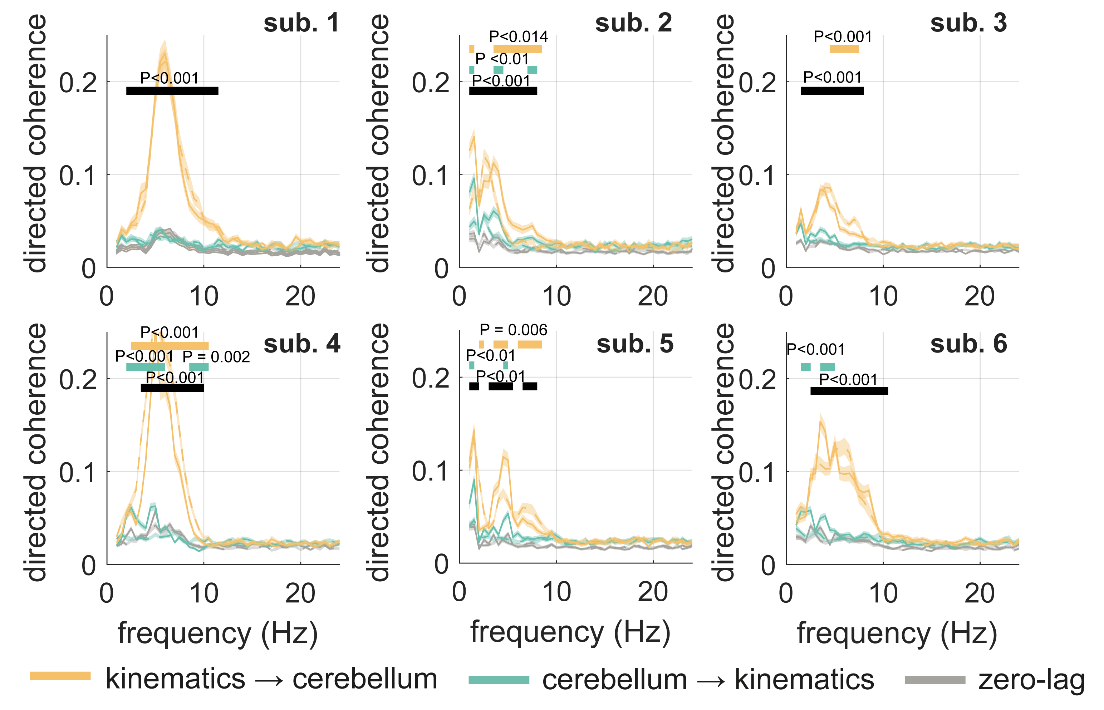

Supplementary Figure 5 - **Summary of changes in cerebellar-accelerometer directed functional connectivity as measured using non-parametric directionality.** Connectivity analyses were made using virtual channels constructed from the ipsilateral cerebellum. Line plots show trial/subject mean, and shaded region gives the standard error of the mean. Directed coherence spectra comparing afferent (kinematics to cerebellum; yellow) and efferent (cerebellum to kinematics; green) showed for each of the six subjects. Bars indicate significant permutation cluster corrected t-statistic for directional connections vs zero-lag (grey).

### 1.7 Supplementary Figure 4

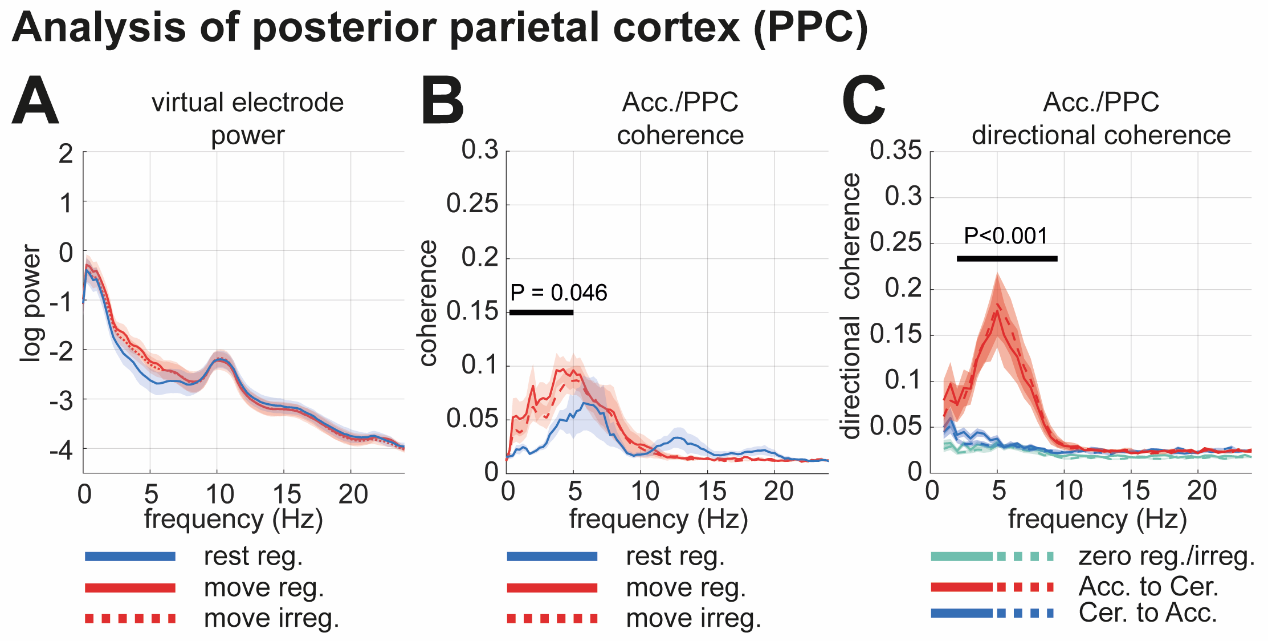

Supplementary Figure 4 – **Analysis of virtual electrode signals projected from a source in the posterior parietal cortex (PPC).** Bars indicate significant cluster permutation statistic (t-test) between rest and movement conditions.

**(A)** Spectral power changes.

**(B)** Coherence between the accelerometry and PPC. **(C)** Directional coherence between accelerometry and PPC estimated using non-parametric directionality.

### 1.8 Supplementary Figure 5

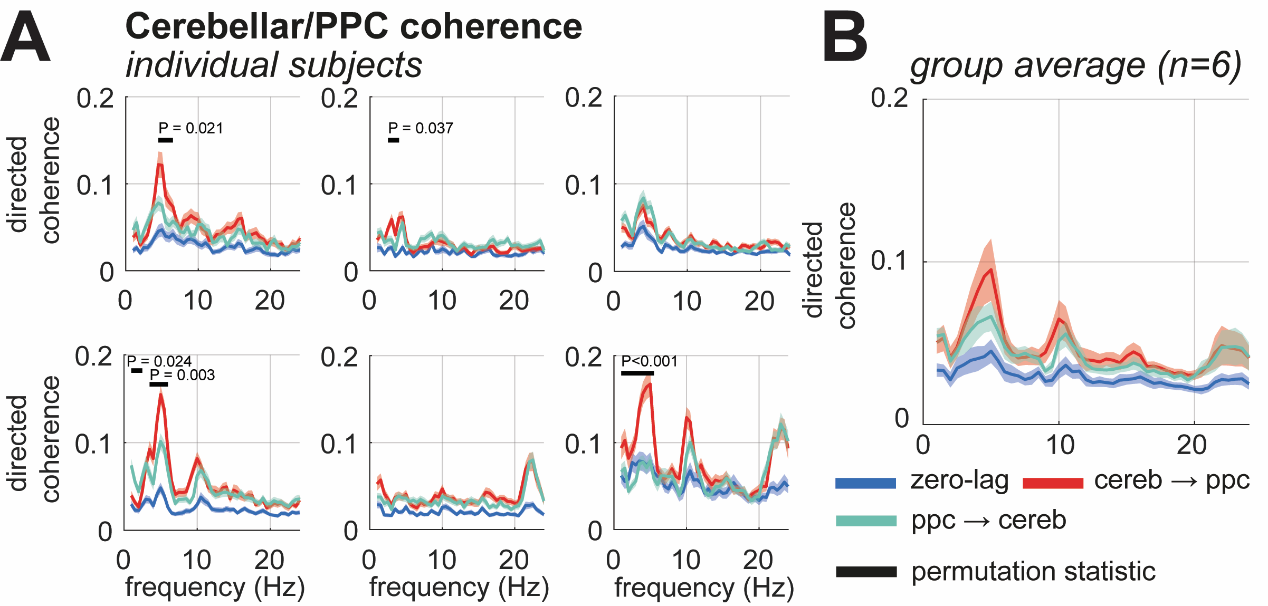

Supplementary Figure 5 – **Analysis of cortico-cerebellar directed functional connectivity.** Estimates were made using non-parametric directionality which accounts for zero-lag shared variance between source localized signals. Bars indicate significant cluster permutation statistic (t-test) comparing forward and reverse directions.

**(A)** Subject level results.

**(B)** Group averaged results.

### 1.9 Supplementary Figure 6

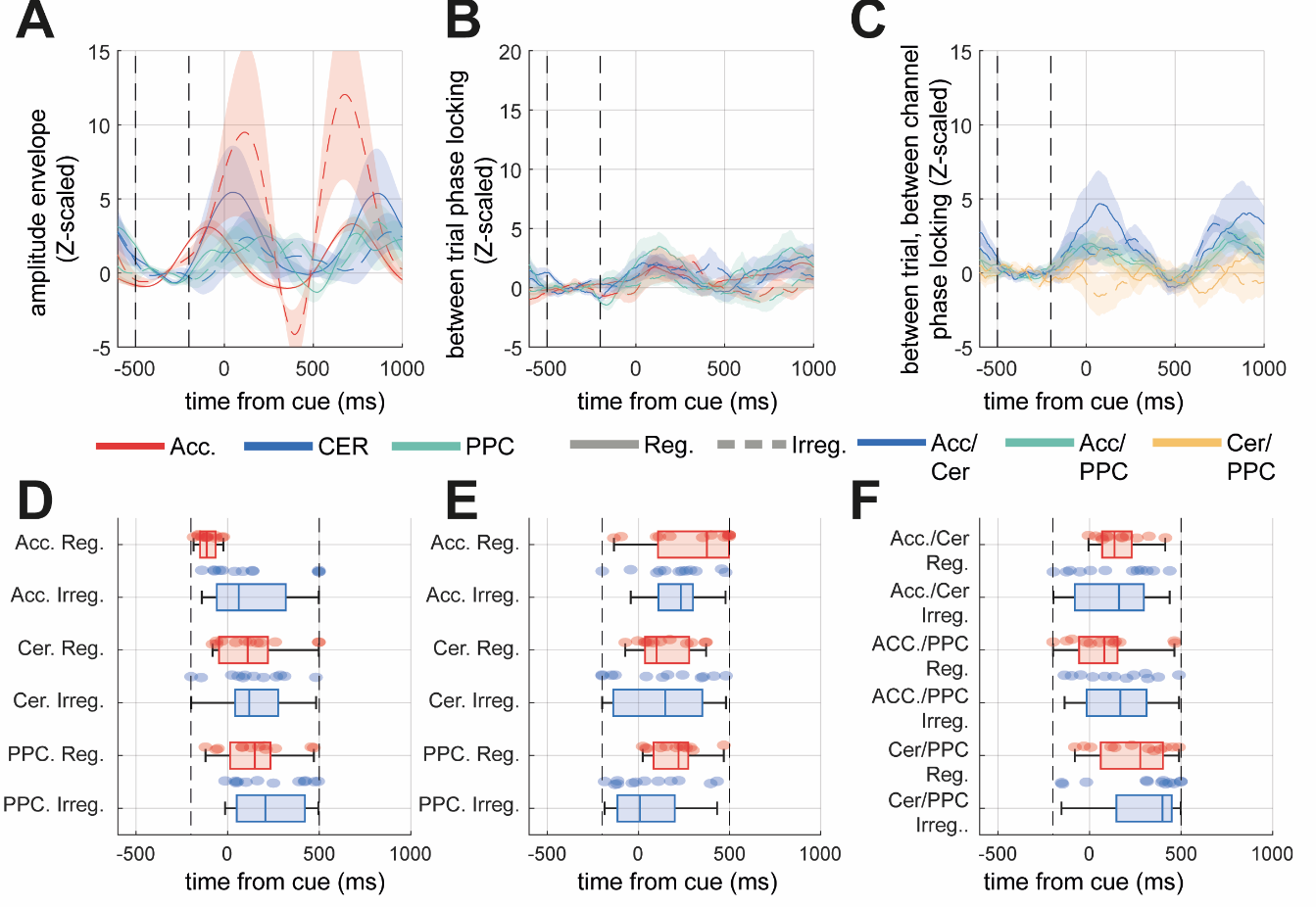

Supplementary Figure 6 – **Cue-locked analysis of** **oscillatory synchronization of cerebellar, posterior parietal cortex (PPC) virtual electrode at frequency of the motor intermittency.** Signals from accelerometry and OP-MEG virtual channels in the cerebellum and posterior parietal cortex (PPC) were time-locked to the onset of the auditory cue (i.e., t = 0).

**(A)** Plot of the group averaged amplitude envelopes of the subject specific motor intermittency band for intermittent acceleration (i.e., accelerometer axis with most power in 4-10 Hz band; red), the motor cerebellum (blue) and the PPC (green).

**(B and C)** Same as A, but for the between trial between-trial phase locking value (PLV, i.e., across trial phase alignment), and between trial, between channel PLV.

**(D-F)** Box plots of time of amplitude envelope peak. Bars indicate P-values of a two-sample t-test exceeding α = 0.05. Dashed lines indicate the ±1 std of the surrogate distribution.

**(E and F)** Same as D and B, but for the between-trial, between-channel PLV (i.e., trial consistency of phase differences), for pairs: Accelerometry/Cerebellum, Accelerometry/PPC, Cerebellum/PPC .

### 1.10 Supplementary Figure 7

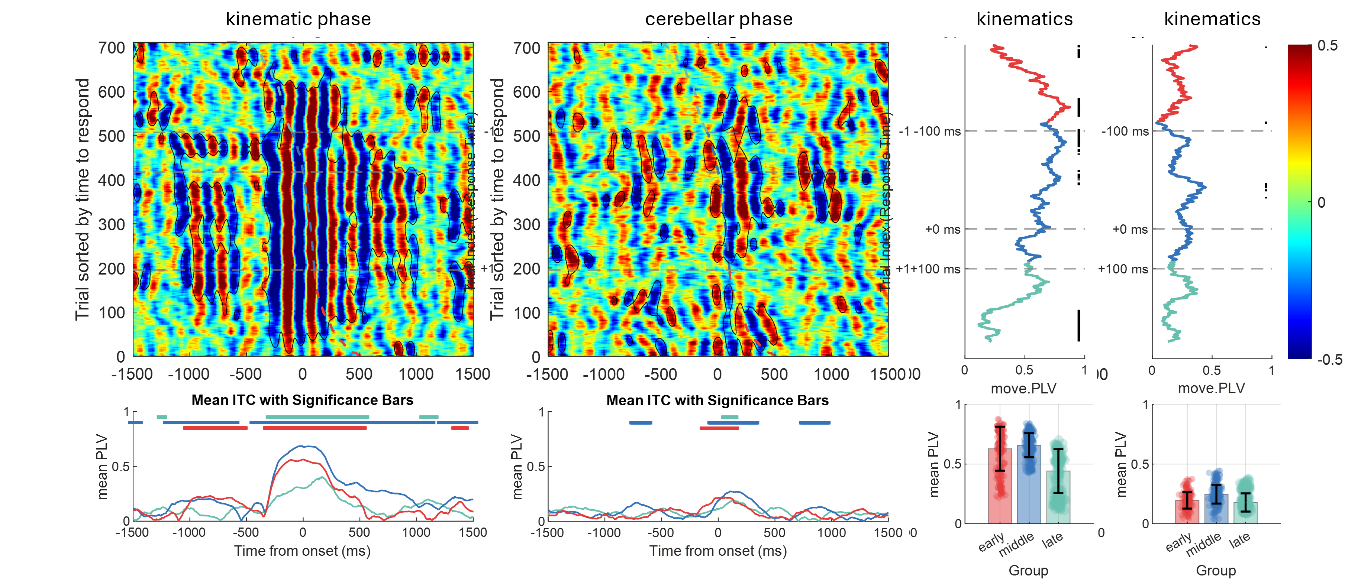

Supplementary Figure 7 – **Example of inter-trial phase synchronization for a single subject.** Instantaneous phase was extracted on a trial-by-trial basis from accelerometery and cerebellar virtual channels, time-locked to peak finger acceleration (t = 0). Trials were categorized by movement timing relative to cue into four groups: early (–500 to –100 ms), anticipatory (–100 to 0 ms), reactive (0 to 100 ms), and late (100 to 500 ms).
